# Placental defects revealed by modelling PWS in mice

**DOI:** 10.64898/2026.01.20.692090

**Authors:** Anna E. Webberley, Raquel Boque-Sastre, Lauren Bailey, Cerys Charles, Rachel A. Jones, Rosie Bunton-Stasyshyn, Michelle Stewart, Sara E. Wells, David S. Chatelet, Stephanie K Robinson, Matthew J. Higgs, Rosalind M. John, Anthony R. Isles

## Abstract

The neurodevelopmental disorder Prader-Willi syndrome (PWS) is caused by loss of paternally-derived gene expression from the imprinted interval on chromosome 15q11-q13. Recently, it has been suggested that the abnormal feeding-related behaviours characteristic of PWS may, in part, be developmentally programmed *in utero* via abnormal placental function. Here we report that several PWS-genes are expressed in mouse placenta with three PWS-transcripts *Magel2*, *Necdin* and the lncRNA *Sngh14*, co-localising to the *Kdr*-positive fetal endothelial cells of the labyrinth zone central to nutrient transport. In a novel PWS deletion mouse model (Large^+/−^) we find markedly reduced expression of PWS genes in the placenta and an associated ∼25% reduction in *Kdr*-positive fetal endothelial cells. Although this did not directly translate into a significant reduction in fetal growth late in gestation, these data suggest that placental function and nutrient transfer from mother to fetus could be compromised in PWS contributing to later post-natal phenotypes.

**Summary statement:** Reduced expression of PWS-associated genes in the mouse placenta results in a 25% loss of the fetal endothelial cells that play a central role in nutrient and gas exchange.

## INTRODUCTION

Prader-Willi syndrome (PWS) is a neurodevelopmental disorder caused by the loss of paternally-derived gene expression from the imprinted cluster on chromosome 15q11-q13 (1). Loss of paternal gene expression primarily occurs because of large deletions spanning the imprinted interval (approximately 70% of cases) but can also result from maternal uniparental disomy (mUPD) or mutations that affect the imprinting control region (ICR) (1, 2). Together, these chromosomal abnormalities and mutations are causal of the majority of PWS cases and lead to complete loss of activity from all paternally expressed imprinted genes within the interval. However, there are also rare micro-deletion cases that highlight certain genes within the interval as being central to the aetiology of PWS (3–5).

Unsurprisingly given the neurodevelopmental nature of PWS, all the paternally expressed genes within 15q11-q13 show high levels of expression in the brain and much of the mechanistic research has focused on neural function in rodent and cell culture models (6–9). Recently however, as well as the direct effects on brain;, there has been a recognition that loss of paternal gene expression may also alter other physiologies which in turn impact brain development and function. One hypothesis is that the metabolic and feeding problems seen in PWS may, in part, be programmed *in utero* via abnormal placental signalling (10). This fits with recent neuroimaging studies showing that structural differences in the hypothalamus of individuals with PWS are consistent across age and distinct from non-PWS obese controls (who show no structural changes), suggesting a developmental origin for these brain changes (11). Interestingly, a role for PWS genes *in utero* has been predicted on the basis of prenatal growth restriction, with the pattern of change indicating constraint of nutrient flow in the later stages of pregnancy (10, 12).

Developmental programming by maternal nutrient provision and/or placental endocrine signalling *in utero* is recognised to impact offspring brain (13, 14). Of relevance to PWS, the development of the melanocortin system, which is a key component of the neural control of feeding and energy homeostasis, is sensitive to changes in a multitude of endocrine and nutritional factors (15). The development and connectivity of the primary constituents of the melanocortin system, pro-opiomelanocortin (POMC) and agouti-related peptide (AgRP) neurons in the hypothalamus and hindbrain, are altered in response to nutrient supply from the mother both pre-natally (via the placenta) (16) and in the early postnatal period (15). For instance, maternal obesity during pregnancy causes a reduction in proliferation of neural precursor cells in the fetal hypothalamus at embryo day 13, which in turn leads to reduced POMC positive neurons and an altered hypothalamic neuropeptide profile and feeding behaviour in adult offspring (17). The placenta directly promotes and protects fetal brain development, buffering the effects of insults such as altered nutrient supply (18, 19). Consequently, it is not unreasonable to assume that loss of gene expression leading to abnormalities in PWS placental function may affect nutrient supply and/or signalling, and could be an additional factor shaping brain development and later life feeding behaviour.

A role for PWS genes in placental function would not be unexpected as the expression of imprinted genes generally are enriched in the mouse (20) and human placenta (21), and genomic imprinting is thought to be intrinsically linked to the evolution of viviparity in mammals (22). Indeed, in early papers describing their discovery, some PWS genes have been shown to be expressed in the placenta (23, 24). However, there has never been a focused study of PWS gene expression in the mouse placenta. The mature mouse placenta, which develops from mid-gestation, is composed of three major layers: the materal decidua and two fetally derived structures, the junctional zone which has a major endocrine function and the labyrinth zone which plays a major role in placental transport (25). Here, we examine PWS gene expression in the placenta across different embryonic timepoints. We show that most PWS genes display imprinted expression in the placenta and also examine the spatial distribution of the three PWS genes: *Magel2*, *Necdin (Ndn)* and the non-coding RNA (lncRNA) *Sngh14* which includes the snoRNAs *Snord115* and *Snord116* (26). All three were localised to the labyrinth zone and fetal endothelial cells in particular. In the Large^+/−^ PWS mouse model, which has a 3.08 Mb deletion encompassing all the paternally expressed genes in this locus (27), loss of expression of PWS genes impacted the cellular composition of the fetal placenta, which showed a reduction in labyrinthine endothelial cells.

Together, these data suggest that loss of gene expression in the placenta of individuals with PWS may have consequences for nutrient transfer across the placenta and raises the possibility of developmental programming of the developing PWS brain.

## RESULTS

### Localisation of *Magel2* to fetal endothelium of the labyrinth

Our preliminary study focused on the PWS and Schaaf-Yang syndrome (28) gene, *Magel2,* which was previously reported to be expressed in the junctional zone of the placenta (29). We initially performed RNAscope *in situ* hybridisation on e14.5 midline sections of wildtype placenta with *Magel2* and *Protocadherin 12* (*Pcdh12)* which identifies glycogen cells located within the junctional zone (Figure 1A) (12). Unexpectedly, the *Magel2* signal did not localise to the junctional zone. Instead, *Magel2* signal specifically localised to the labyrinth. To further define the localisation of *Magel2*, we co-hybridised *Magel2* with markers that distinguish cell types in the labyrinth using *Cathepsin Q* (*Ctsq)* for the sinusoidal trophoblast giant cells and *Kinase Insert Domain Receptor* (*Kdr*; *Flk1*) for the fetal endothelium. *Magel2* signal co-localised exclusively with *Kdr-*positive cells indicating expression in the fetal endothelium of the placental labyrinth (Figure 1B).

**Figure 1.**
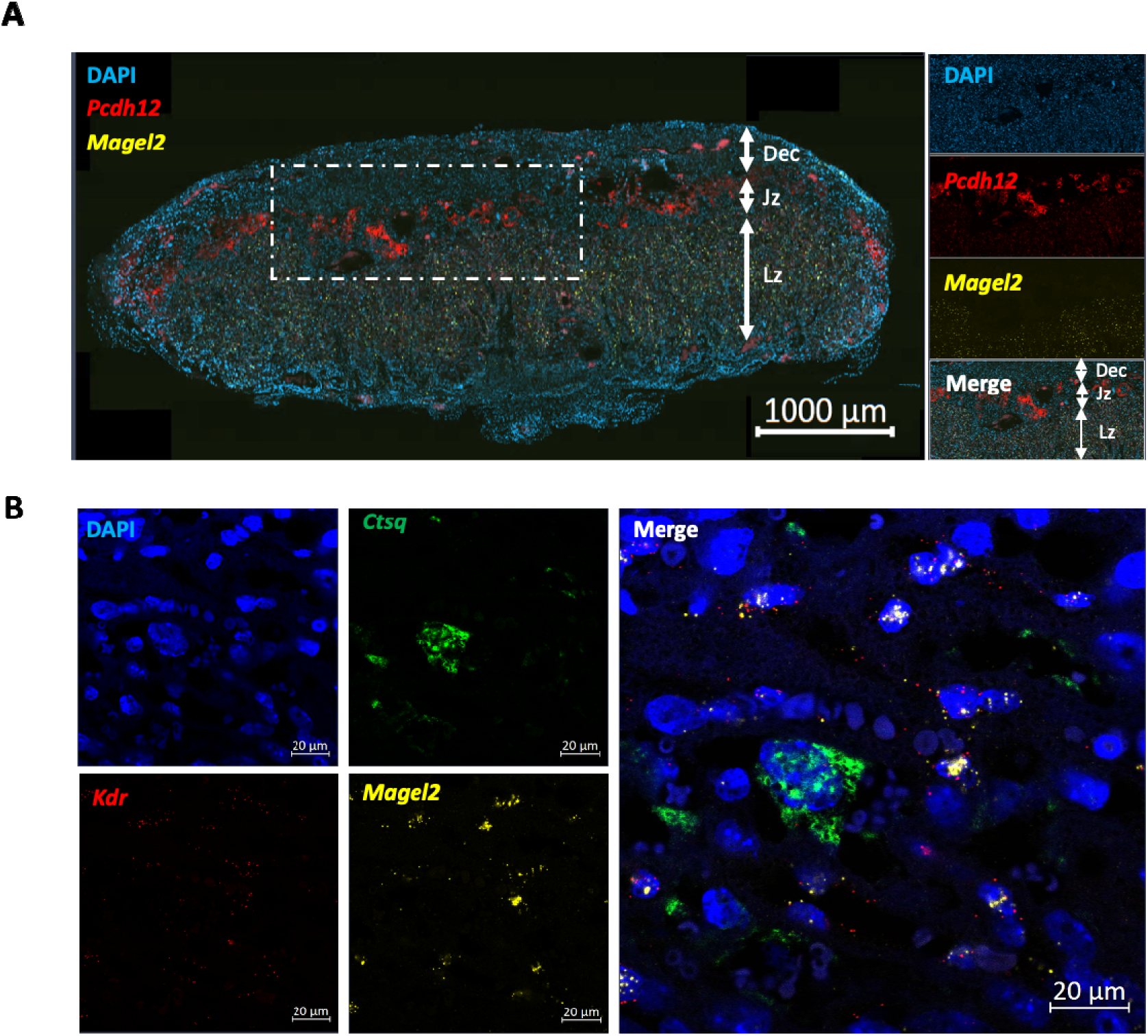
*Magel2* localisation to the fetal endothelium of the mouse placenta. Midline section of e14.5 mouse placenta hybridised with RNAscope probes to (A) *Pcdh12* (glycogen cells located in the junctional zone) and *Magel2* or (B) *Ctsq* (sinusoidal cells of the labyrinth) and *Kdr* (fetal endothelial cells of the labyrinth) and *Magel2* localised *Magel2* to the fetal endothelium. DAPI = 4′,6-diamidino-2-phenylindole; Dec = maternal decidua; Jz = junctional zone; Lz = labyrinth zone.

### Multiple PWS genes are expressed in mouse placenta

We then expanded our assessment of PWS genes in the placenta. The expression level of six genes was examined in wild-type whole placenta by RT-qPCR at e10.5 and e12.5, when the mature placenta is rapidly expanding, and at e16.5 which is the time of maximum placental growth (12). We chose to analyse the expression of the three protein coding genes, *Necdin (Ndn), Mkrn3* and *Magel2,* and three representatives of individual transcripts from lncRNA *Snhg14,* specifically, the snoRNAs *Snord115* and *Snord116* and the ncRNA *Ipw.* The protein coding genes were chosen as they have tractable cellular functions, and the *Snhg14* transcripts on the basis that they are critical to the incidence of the core PWS phenotype (5, 30), and that *Ipw* is a regulator of the imprinted Dlk1-DIO3 locus (31), which has a known role in placental function (32).

Levels of expression varied between the different genes, with the highest being *Ndn, Snord115* and *Snord116,* the lowest being *Ipw*, with *Mkrn3* and *Magel2* being intermediate (Figure 2). The Angelman syndrome gene, *Ube3a*, also showed robust expression in wild-type placenta (Figure S1A). Whilst *Mkrn3* showed relatively stable expression across all three timepoints (no main effect of TIMEPOINT, χ^2^ statistic=1.51, *P*=0.25), both *Ndn* (main effect of TIMEPOINT, χ^2^ statistic=8.66, *P*=0.013) and *Ipw* (main effect of TIMEPOINT, χ^2^ statistic = 3.93, *p* = 0.04) decreased in placental expression with increasing embryonic age. *Post hoc* testing (Dunn pairwise) revealed that for *Ndn* this change was driven by a significant decrease in expression between e12.5 and e16.5 (Z statistic = 2.86, Adj. *p* = 0.004), whereas *Ipw* decreased between e10.5 and e12.5, although this change did not survive Bonferroni correction (Z statistic = 2.19, Adj. *p* = 0.09). In contrast, *Magel2* placental expression increased across embryonic timepoints (main effect of TIMEPOINT, χ^2^ statistic = 15.86, *p* = 0.0004), with a significant increase between e10.5 and e12.5 (Z statistic = –3.75, Adj. *p* = 0.0005). The snoRNAs demonstrated a pattern of expression similar to *Ndn*, with both *Snord115* (Z statistic = 3.01, Adj. *p* = 0.008) and *Snord116* (Z statistic = 2.58, Adj. *p* = 0.03) decreasing between e12.5 and e16.5. However, both had highly variable levels of expression with *Snord115* (χ^2^ statistic = 5.83, *p* = 0.011), but not *Snord116* (χ^2^ statistic = 3.07, *p* = 0.06) demonstrating a main effect of TIMEPOINT overall.

**Figure 2.**
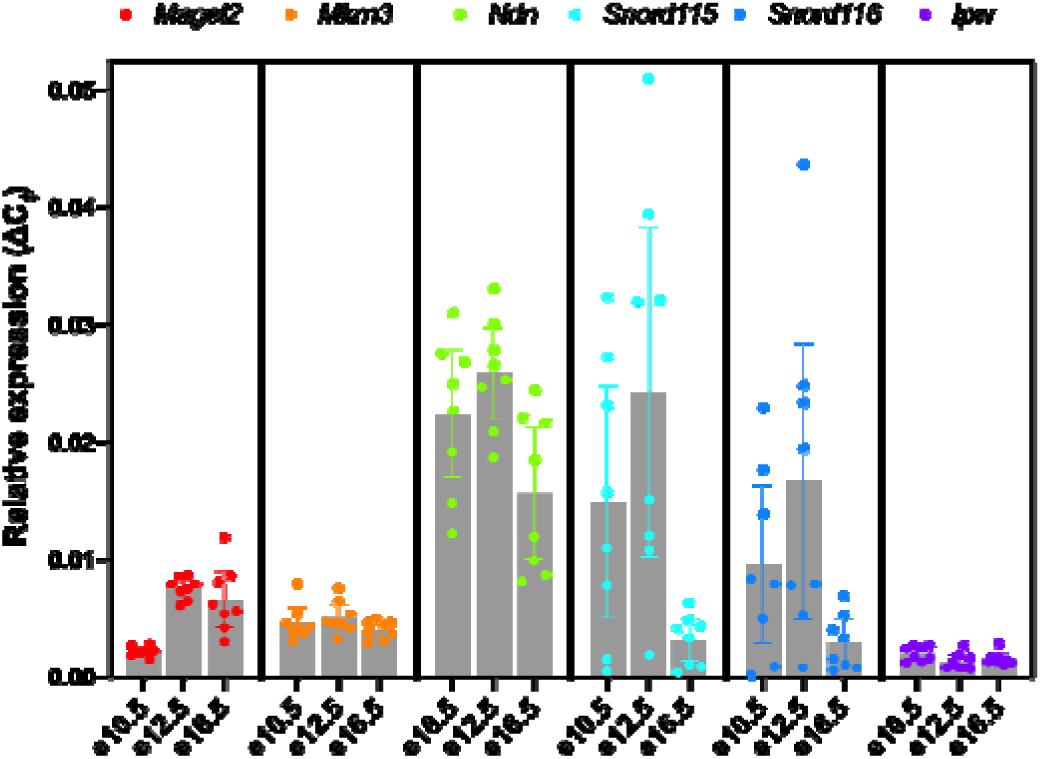
Expression of PWS genes in wild-type placenta homogenate at three different embryonic timepoints (e10.5, e12.5 and e16.5). Gene expression normalised to the geometric mean of three house-keeping genes using the ΔCt method. N=8 (4M,4F) for each timepoint. Bars are means with 95% CI indicated.

No gene expression data demonstrated a main effect of SEX, or an interaction between TIMELINE and SEX. Full statistical details can be found in Tables S1-S3 (Supporting Information).

### Reduction in placental PWS gene expression in Large^+/−^ mice

Some imprinted genes demonstrate tissue specific imprinting (21). We took advantage of the Large PWS mouse model carrying 3.08 Mb paternal deletion (Large^+/−^) encompassing all the genes in the locus (27) to determine how placental expression of these genes was altered in response to a common PWS mutation. Expression of the same six PWS genes in placenta from Large^+/−^ and wild-type littermates was examined using RT-qPCR at two embryonic timepoints, e12.5 and e16.5. At e12.5 (Figure 3A), expression of all PWS genes was significantly reduced in Large^+/−^ mice following correction for multiple testing (see Table S4 and S5 for full statistics). At e16.5, the picture was broadly the same, with all PWS genes showing a decrease in expression in Large^+/−^ placenta compared to wild-type (Figure 3B).

**Figure 3.**
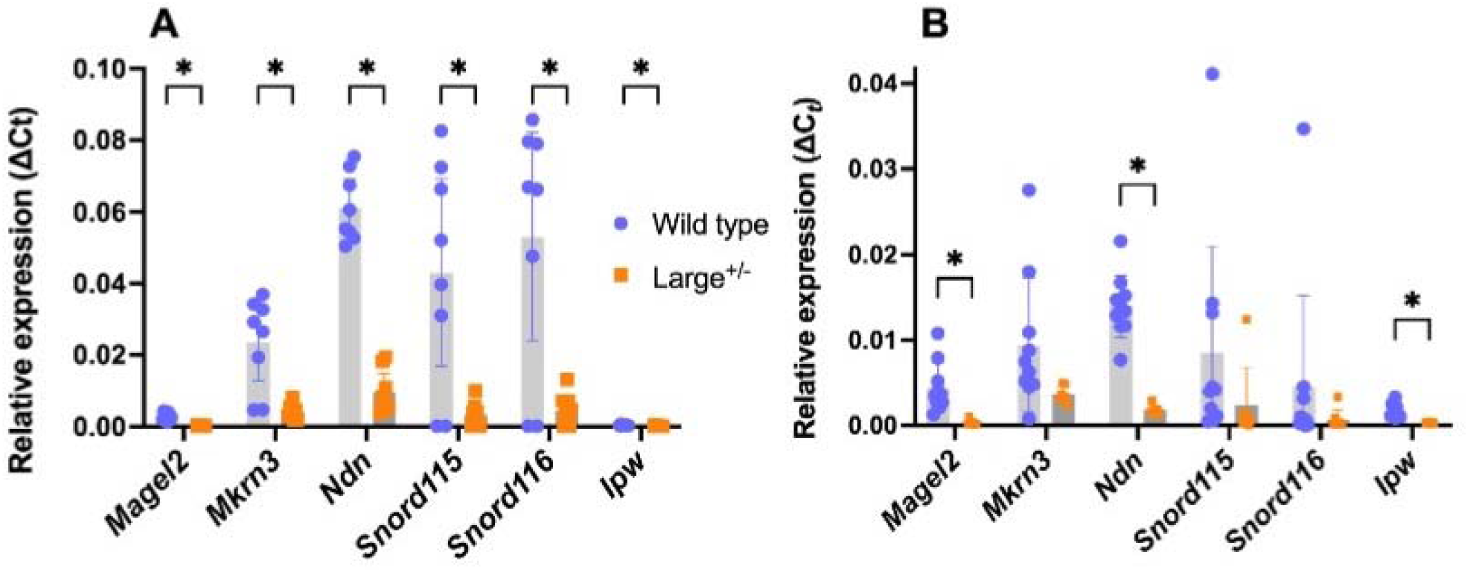
Expression of PWS genes in placenta homogenate from wild-type and Large^+/−^ mice at two different embryonic timepoints. At e12.5 (**A**), expression of all PWS genes in the placenta was significantly reduced in Large^+/−^ mice relative to wild-type controls. At e16.5 (**B**), although all average expression was reduced in Large^+/−^ mice, the data were quite variable, and this difference only reached statistical significance for three PWS genes. Gene expression normalised to the geometric mean of three house-keeping genes using the ΔCt method. N=8 (4M,4F) for each genotype at both timepoints. Based on Shapiro-Wilk test, normally distributed data were analysed by ANOVA using the factors GENOTYPE and SEX; non-normally distributed data as determined by the Shapiro-Wilk test, were analysed by Kruskal-Wallis using the factors GENOTYPE and SEX. P values indicated on the graphs are adjusted to account for multiple testing using False Discovery Rate (FDR) correction. Bars are means with 95% CI indicated.

However, only three of these reached statistical significance following correction for multiple testing (see Table S6 for full statistics). There was no difference in expression of the maternally expressed Angelman syndrome *Ube3a* at e12.5 and or e16.5 (Figure S1A). There was no difference in *Ube3a* expression between males and females, and no significant interactions between GENOTYPE and SEX at either timepoint.

This pattern of reduced PWS gene expression matches expression in Large^+/−^ fetal brain relative to wild-type at the two timepoints (n=4 per genotype, 2M and 2F; Figure S2). The same is true for *Ube3a*, that showed robust expression, equivalent to wild-type in Large^+/−^ fetal brain (Figure S1B).

### Localisation of *Ndn* and *Snhg14* to the fetal endothelium of the labyrinth

We next asked whether other PWS genes were expressed in the fetal endothelium focusing on the most highly expressed PWS genes notably altered in the PWS model, *Ndn, Snord115* and *Snord116*. Due to technical difficulties designing probes specifically to *Snord115* and *Snord116,* we used an off-the shelf probe for *Snhg14* which is the lncRNA encompassing *Snord115*, *Snord116* and *Ipw* (26). In the wild-type placenta expression of both *Ndn* and *Snhg14* was localised to the labyrinth zone of the placenta (Figure 4A-D) with ∼20% of wild-type placental cells expressed *Ndn* and ∼30% expressed *Snhg14* (Figure 4A-D). Less than 1% of placental cells in the Large^+/−^ mouse expressed these same genes (Figure 4E. Mann-Whitney, *Ndn p* = 0.0002; *Sngh14 p* = 0.0002). More detailed analysis of expression in the labyrinth demonstrated that 51% of *Ndn* expressing, and 47% *Snhg14* expressing cells also expressed the fetal endothelium marker *Kdr* (Figure 4C and D). We further tested this by counting the coincidence of *Ndn* and *Snhg14* molecules with *Kdr*-positive and *Kdr*-negative cells. The average number of molecules of *Ndn* and *Sngh14* was significantly higher in *Kdr*-positive cells than that of *Kdr*-negative cells (Figure 4F, Wilcoxon matched-pairs signed rank test, *P* = 0.7891 for both *Ndn* and *Sngh14*).

**Figure 4.**
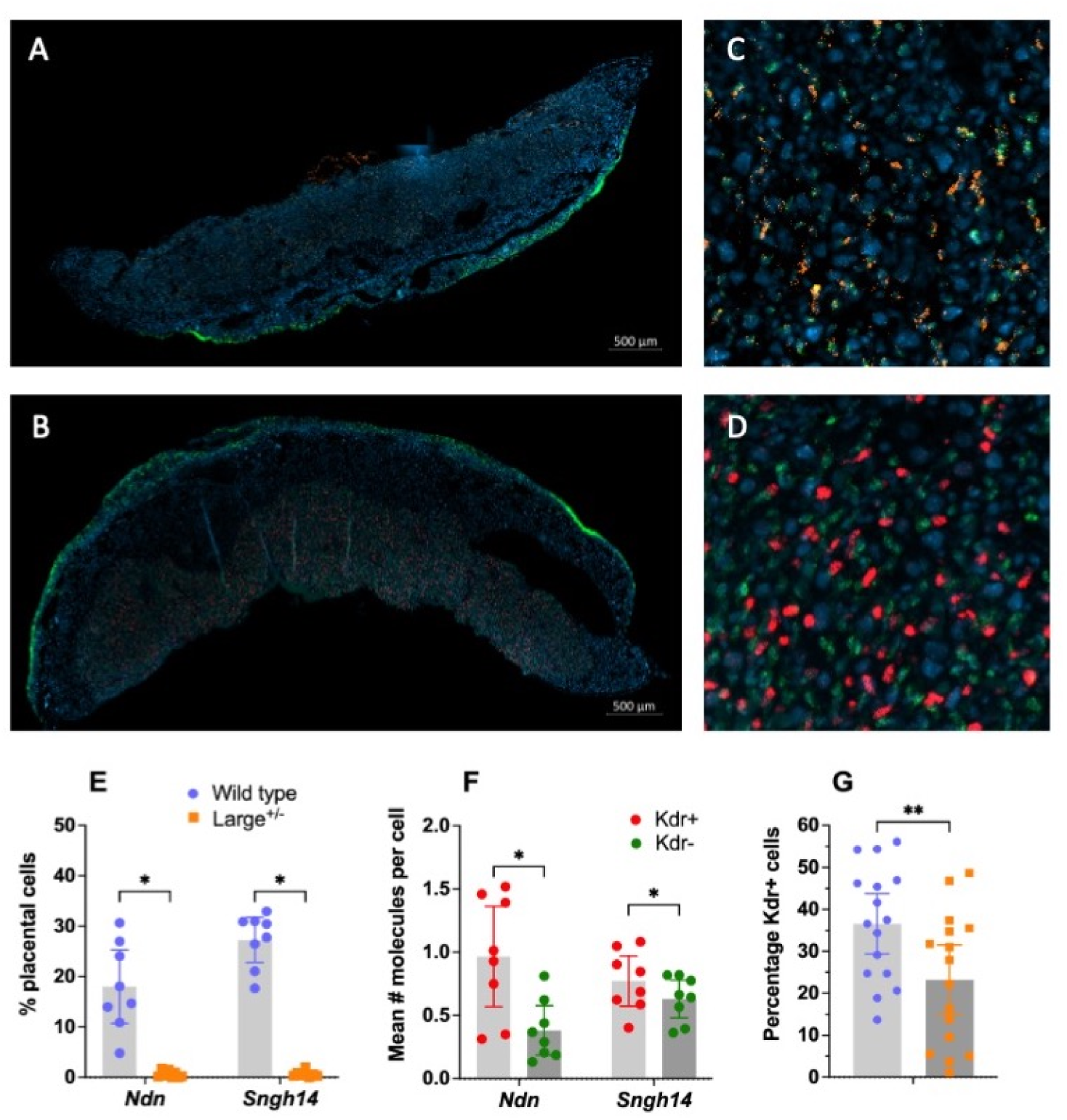
RNAscope analysis of *Ndn* and *Snhg14* gene expression in endothelial cells of labyrinth zone of the placenta. Representative image of RNAscope analysis of *Ndn* (orange) (**A**) and *Snhg14* (red) (**B**) indicated strong expression of both genes in the labyrinth zone of the wild-type placenta, with a high degree of co-localisation with the endothelial cell marker *Kdr* (green) (**C** – *Ndn*; **D** – *Snhg14*). Proportion of all placental cells expressing *Ndn* and *Snhg14* in wild-type and Large^+/−^ placenta (Mann-Whitney) (**E**). Quantification of co-localisation indicated that both genes were statistically (Wilcoxon matched-pairs signed rank test) more likely to be expressed in a *Kdr*+ cell than a *Kdr-* one (**F**). Quantification of total cell number and *Kdr*+ cells in demonstrates a reduction (Wilcoxon matched-pairs signed rank test) in the proportion of *Kdr*+ cells in Large^+/−^mouse relative to wild-type littermates (**G**). Bars are means with 95% CI indicated.

### Loss of endothelial cells in the labyrinth in Large+/− placentae

We then went on to examine if loss of expression of these and other PWS genes in Large^+/−^ placenta was associated with changes in the number of fetal endothelial cells. There was a significant reduction in the proportion of *Kdr*-positive cells in the labyrinth of Large^+/−^ placenta compared to wildtype (Figure 4G, ANOVA F= 6.878, P=0.014).

### No gross difference in Large^+/−^ placentae size

Placental midline sections from Large^+/−^ (n=10; 6M, 4F) and WT (n=13; 7M, 6F) were H&E stained (n=4 per sample) and used to determine the midline area. There were no differences in area between Wild-type and Large^+/−^ mice (Table 1).

**Table 1.**
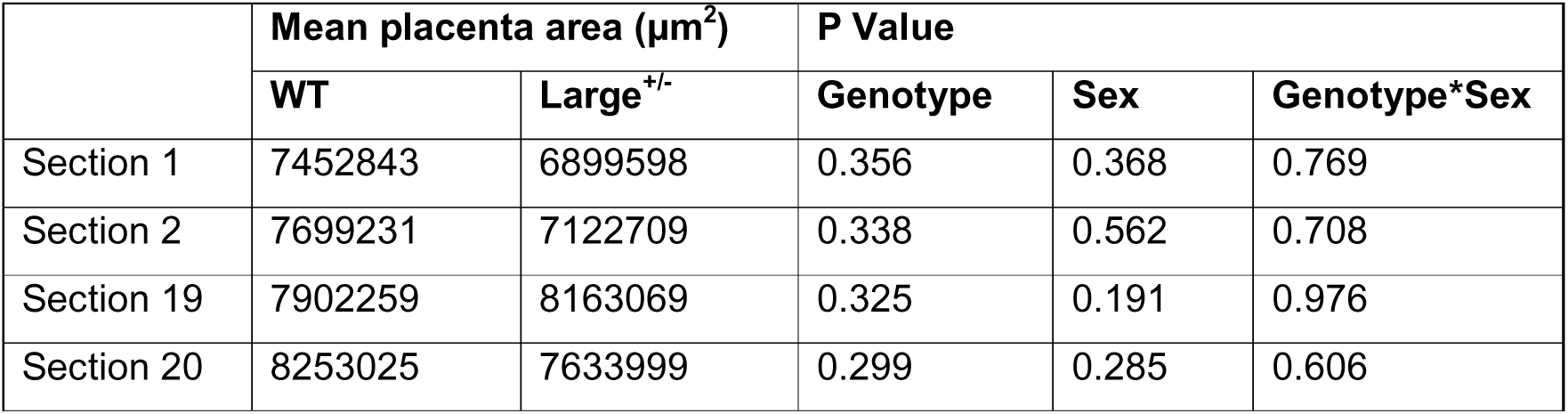
Area measurements for placental sections. The area of equivalent placentae sections were measured. A Levene’s test was used to determine normality of distribution (significance threshold P=0.05). As all data was found to be normally distributed, a two-way ANOVA was used to determine if there was a significant difference in area between GENOTYPE, SEX or if there was a GENOTYPE*SEX interaction. An initial significance threshold of P=0.05 was used, which following correction for multiple testing using the Bonferroni method was reduced to P=0.0125.

### Fetal development in Large^+/−^ mice

Large^+/−^ mice show almost complete lethality at birth (27), and so the potential impact of abnormal placenta signalling on Large^+/−^ fetal growth was assessed at E18.5 using MicroCT. Although the volumes of whole embryos, kidney and brain were generally lower for Large^+/−^ mice compared to wild-type, none of these reached statistical significance (Table 2). The average intensity (mean brightness [lower, upper 95% CI]) of Large^+/−^ kidney (35132 [32348, 37917]) and brain (26095 [23189, 29001]), were not significantly different from wild-type kidney (33714 [30929, 36498], F_1,9_=0.66, P=0.24) or brain (26089 [23182, 28995], F_1,9_=0.001, P=0.997).

**Table 2.** Whole embryo and selected tissue volumes in E18.5 Large^+/−^. Reductions in whole embryo, and kidney and brain volumes were seen in Large^+/−^ fetuses but none of these reached statistical significance in comparison to wild-type.

|  | Mean (95% confidence interval) |  | F<sub>1,9</sub> | P | Partial η<sup>2</sup> |
| --- | --- | --- | --- | --- | --- |
|  | Wild-type (n=6; 3F, 3M) | Large<sup>+/-</sup> (n=6; 3F, 3M) |  |  |  |
| **Whole Embryo** | 1.11cm<sup>3</sup> (0.94,1.28) | 0.91cm<sup>3</sup> (0.74, 1.08) | 3.36 | 0.100 | 0.272 |
| **Kidney** | 7.83e+9μm<sup>3</sup><br>(6.44e+9, 9.21e+9) | 6.80e+9μm<sup>3</sup><br>(5.42e+9, 8.19e+9) | 1.40 | 0.266 | 0.135 |
| **Brain** | 7.63e+10μm<sup>3</sup><br>(6.92e+10, 8.34e+10) | 6.99e+10μm<sup>3</sup><br>(6.28e+10, 7.70e+10) | 2.12 | 0.179 | 0.191 |

## DISCUSSION

The idea that changes in nutrient supply and/or signalling from the placenta during development may be an additional impact on PWS brain development and function is gaining traction (10, 11). Here, we explored PWS gene expression in the mouse placenta and the consequences of loss of this expression in the Large^+/−^ mouse line, which models the genetic lesion seen in approximately 70% of PWS cases (1, 2). We demonstrated that many PWS genes are expressed in the placenta, specifically the fetal endothelium, and that expression varies across different developmental stages. Loss of expression of PWS genes in the Large^+/−^ model (27) impacted the cellular composition of the labyrinth zone of the placenta, which showed a reduction in fetal endothelial cell number. Although no detectable difference was seen in Large^+/−^ fetal size late in gestation, these data suggest that loss of gene expression in the placenta of individuals with PWS could potentially have functional consequences, highlighting the need for further pre-natal studies.

The role of the placenta changes across development with expansion of the endocrine lineages from e9.5-e16.5 driving maternal adaptations and the labyrinth continuing to expand until term to support substantial fetal growth (12). We examined expression at key timepoints, capturing when the mature placenta is rapidly expanding (e10.5 and e12.5) and the time of maximum placental growth (e16.5), to identify whether PWS gene expression also changed across this period. *Mkrn3* and the non-coding RNA *Ipw* both showed little change across the time points examined, but these two genes were also the lowest expressed transcripts. The highest expressed PWS transcripts, *Ndn, Snord115* and *Snord116*, all had peak expression levels at e10.5 and e12.5, decreasing significantly at e16.5. In contrast, another robustly expressed PWS gene, *Magel2*, demonstrated a significant increase in expression between e10.5 and e12.5, which was maintained at e16.5. These expression patterns across embryonic time points, relative levels and inferred spatial location can also be seen when exploring recent mouse placenta scRNA-seq database that spans mid-gestation (e7.5 to e14.5) (33). In the placenta of the Large^+/−^ mouse model, which has a 3.08 Mb deletion encompassing all the paternally expressed genes in this locus (27), PWS gene expression was reduced at the two time points assessed, indicating that these genes show parent-of-origin specific expression, at least in a majority of cells in the placenta with potential to contribute to placental dysfunction in PWS.

When we explored the spatial distribution of a subset of PWS transcripts. *Magel2*, previously thought to be expressed in the junctional zone (29), was instead co-localised with *Kdr+* fetal endothelial cells in the labyrinth. Similarly, the two most highly expressed PWS genes, *Ndn* and the lncRNA *Sngh14,* which includes the snoRNAs *Snord115* and *Snord116* (26), were also highly expressed in *Kdr+* fetal endothelial cells, both in terms of their coincidence and expression levels. A number of *Kdr*-negative cells in the labyrinth also expressed *Snhg14* but expression levels of *Ndn* and *Snhg14* were significantly higher in *Kdr*-positive cells. Together, these data support a non-random distribution of PWS gene expression in the labyrinth zone.

Although there was no gross change in overall placental morphology detected, counting of *Kdr*-expressing endothelial cells revealed that there was a reduction of approximately 25% in the labyrinth zone of Large^+/−^ mice relative to wild-type controls. This is of interest because *Dlk1*, another paternally expressed imprinted gene, localises to the same cell type (34) further hinting at convergence of imprinted function on this cell type.

The labyrinth zone is responsible for nutrient and oxygen exchange between mother and foetus (35). In particular, the endothelial cells of the labyrinth zone play an important role in vasculogenesis, angiogenesis and maintenance of vasodilation in existing blood vessels (36, 37). Although placental vascularisation was not measured directly here, changes can have an impact blood flow throughout the placenta which, in turn, is vital for nutrient and oxygen transfer between mother and fetus (38). Small size has been reported as a feature of Prader-Willi syndrome (39–42) and the pattern of effects could reflect nutrient restriction in the later stages of pregnancy when large amounts of fat are deposited, fitting with the idea of maternal and paternal imprinted genes being in conflict over resources (43). Using MicroCT on a small number of fetuses we were not able to detect a difference in Large^+/−^ fetal size at e18.5, although we recognise that studies with larger numbers are required. Interestingly, in the PWS imprinting centre deletion mouse model in which all PWS gene expression is also lost, surviving pups are leaner from birth (44).

A reduction in nutrient supply because of reduced endothelial cells could lead to abnormal programming of the fetus, even in the absence of clear effects on fetal growth (38). Certainly, developmental programming by maternal nutrient provision *in utero* is recognised to impact offspring brain (13, 14, 45), including the developing melanocortin system which is a key component of the neural control of feeding and energy homeostasis (15). A two-hit hypothesis has been proposed, whereby loss of PWS gene expression has direct consequences for brain development, but also indirectly effects via abnormal placental signalling (10). Our findings that PWS genes are expressed in the placenta and that loss of expression leads to changes in the cellular composition of the labyrinth zone would suggest this is a possibility, and that nutrient transfer across the PWS placenta needs further exploration.

## MATERIALS AND METHODS

### Animals and samples

All animal studies were licensed by the Home Office under the Animals (Scientific Procedures) Act 1986 Amendment Regulations 2012 (SI 4 2012/3039), UK, and additionally approved by the Institutional Ethical Review Committees. All animal studies complied with the ARRIVE guidelines.

Wild-type placenta samples were generated from timed mating of C57BL/6J or 129SV mice (*Magel2 in situ*) within Cardiff University. The morning a plug was detected was recorded as embryo day 0.5 (e0.5). Pregnant females were housed singly in cages in a vivarium maintained on a 12-hour light-dark cycle (lights on from 08:00-20:00) at a temperature of 22 (± 2) °C and 50 (± 10) % humidity. Standard laboratory chow and water was available *ad libitum* throughout all experiments. Samples were collected between 09:00 and 11:00.

The PWS large deletion (C57BL/6J-Pcan^em4H^/H, called Large^+/−^ here) model was generated as part of the Foundation for Prader-Willi Research Pre-Clinical Animal Network (FPWR-PCAN) by MRC Mary Lyon Centre, Harwell Oxford, UK (MRC MLC) as previously described (27). Briefly, the large deletion (3.08 Mb) spans 2820bp upstream of the *Mkrn3* gene and 933bp downstream of *Ube3A*. As such, the deletion removes all the PWS critical genes including *Mkrn3* but excludes maternally expressed *Ube3a*. The Large^+/−^ mice were also maintained on C57BL/6J mice and were produced, genotyped, and sexed at MRC MLC.

Large^+/−^ (4M, 4F) and wild-type (4M, 4F) littermate samples were generated from three separate timed mating of Large^+/−^ positive males with C57BL/6J female mice. The morning a plug was detected was recorded as e0.5. At MRC MLC, pregnant females were singly housed in IVCs (Tecniplast BlueLine 1284), on grade 4 aspen wood chips (Datesand, UK), with shredded paper shaving nesting material and small cardboard play tunnels for enrichment. The mice were kept under controlled light (light, 07:00–19:00; dark, 19:00–07:00), temperature (22 ± 2 °C) and humidity (55 ± 10%) conditions. Standard laboratory chow and water was available *ad libitum* throughout all experiments. Fetal brain and placenta were collected at e12.5 and e16.5 between 09:00 and 11:00. One half of the placenta was fresh frozen on dry-ice and the other fixed in 10% formalin for 24 hours then processed and wax-embedded. Brain samples from embryos were fresh frozen on dry-ice. All samples were shipped to Cardiff University for downstream processing.

Sample size was determined via Power Calculation (46), and was based on difference between controls and experimental groups of 40%, with an overall standard deviation of 25%, power of 0.8 and alpha of 0.05; n=7 is required.

### RNA extraction and RT-PCR

Samples were transferred to lysing matrix tubes with TRI reagent (Zymo research, California) (1000µl for brain samples, 700µl for placenta samples). These tubes were placed in a Fast Prep® FP120 machine for 3 x 10 second long cycles at a speed of 5m/s. Tubes were spun in a centrifuge at room temperature for 5 minutes, at a speed of 13000 rpm, which was used throughout. The supernatant was extracted from the lysing matrix tubes and transferred to an RNase free tube, and an equal volume of ethanol (95-100%) was added and mixed thoroughly. RNA was then extracted using Direct-zol™ miniprep kit as per the manufacturer’s instructions. The RNA was eluted using 50µl of DNase/RNase free water and then stored at – 80°C until needed.

RNA quality and concentration was assessed using a Nanodrop™ 8000 Spectrophotometer (Thermo Fisher Scientific, Massachusetts, USA). Good quality RNA (260/280 result of as close to 2 and >1.8) was converted to cDNA. 2 µg of RNA was diluted in 20ul RNase-free water and added to cDNA EcoDry™ Premix (Double Primed) (Takara/ Clonetech laboratories, California US) and mixed thoroughly with a pipette. These were then subject to PCR under the following conditions: 42°C for 1 hour, 70°C for 10 minutes. Each 20µl sample was then diluted in 180µl of nuclease free water. The samples were then stored at –20°C.

### Quantitative PCR (qPCR)

qPCRs were set up using a Corbett Robotics CAS-1200 (Qiagen, Hilden, Germany). A master mix was made up containing 12.5µl SensiMix SYBR No-ROX Kit (Bioline, London, UK), 1.75µl of forward and reverse primer for the chosen gene, 4µl nuclease free water and 5ul of sample. A no template control (NTC) of nuclease free water was also added for each qPCR run and all samples were run in triplicate. The PCR was run under the following conditions: 95°C for 10 minutes, then 40 cycles of: 95°C for 20 seconds, 60°C for 20 seconds, and then 72°C for 20 seconds. Following amplification a melt curve was generated by increasing the temperature in 1 °C increments between 50°C and 99 °C, and holding each change in temperature for 5 seconds. Sequences for primers used in this study can be seen in Table S7.

qPCR data were processed using the delta *C*_t_ method (Livak and Schmittgen 2001) and for each tested gene the results were normalised to a geometric mean of the housekeeping genes (*Hprt, Dynein* and *B-actin*). The Levene-test was used to test normality of the data. If normally distributed, the data were analysed by two-way ANOVA of the delta *C*_t_ values. For the analysis of wild-type PWS gene expression across development, the factors TIMEPOINT (e10.5, e12.5, e16.5) and SEX (male, female) were applied. To analyse Large^+/−^ data, the factors GENOTYPE (wild-type, Large^+/−^) and SEX (male, female) were used. For non-normally distributed data, a log transformation was first carried out to see if this would cause the data to assume normality, if the data was still not normally distributed then a Kruskall-Walis test was used as a non-parametric equivalent significance test. Both tests used significance thresholds of p<0.05. P values were then adjusted to account for multiple testing using False Discovery Rate (FDR) correction.

### Image analysis of H&E stained placentas

H&E stained placenta sections from wild-type and Large^+/−^ mice were imaged and measured using the Zen image analysis tool. These area values were then analysed in RStudio using a Levene test to check for normality and a two-way ANOVA with the factors GENOTYPE (wild-type, Large^+/−^) and SEX (male, female) to determine if there was a difference in placenta areal. Both tests used initial significance thresholds of p<0.05. The Bonferroni method was used to correct P values for multiple testing before determining if a result was significant.

### RNAscope

RNAscope ® Multiplex Fluorescence Assay was performed according to the manufacturers guidelines essentially as previously described (47) using RNAscope ® Multiplex Fluorescence reagent kits (ACD Bio, Newark, CA, USA). For *Magel2* localisation, e14.5 midline sections of wildtype 129/Sv placenta were hybridised with probes to *Pcdh12* (ACD-Mm-Kdr-489891-C1; 1/1:500) and *Magel2 (*ACD-Mm-Magel2-502971-C3; 1:1000) or probes to *Ctsq* (sinusoidal cells of the labyrinth; ACD-Mm-Ctsq-423681-C1); *Kdr* (fetal endothelial cells of the labyrinth; 414818-C2) and *Magel2*. For quantification of fetal endothelium and localisation of *Ndn* and *Snhg14* wild-type and Large^+/−^ e16.5 BL6 midline placental sections were hybridised with probes to *Kdr* and either *Ndn* (ACD-Mm-Ndn-442711-C3) or *Snhg14* (ACD-Mm-Snhg14-C3). *Snhg14* (the lncRNA containing *Snord115*, *Snord116*, and *Ipw*) was chosen as it was not possible to design probes to the individual sno– and ncRNAs. N=8 placenta of each genotype with two sections counted per placenta. Each slide contained two placenta sections, one was treated with probes for the aforementioned genes (termed the ‘probe section’), one was treated with probe dilutant as a control, to indicate what parts of the image produced were true signal, and what was only background (Figure S3).

### RNAscope Image Analysis

Slides were imaged using the Zeiss Fluorescence microscope, with each channel for each group being set to the same light exposure times. Images were taken within 1 week of the slide being produced. Images were then processed using Zen blue 3.5 imaging software. Background subtraction and Gauss settings were used to pre-process the images, before a background fluorescence value was calculated from an adjacent no-probe control section on the slide. Only signal intensity above this no-probe threshold value was deemed ‘true signal’. The analysis gave the number of positive cells for each gene within each placenta section, and the number of molecules of ‘true signal’ found within each positive cell. The proportion of cells showing true signal within each placenta section for each tested gene was then calculated, as well as the proportion of cells showing colocalization, and the data was compared for wild-types and Large^+/−^ mice. For the percentage of *Kdr* positive (endothelial) cells per placenta, an asin/sqrt transformation was first used, followed by a two-way ANOVA to determine if there was a statistically significant effect of the factors GENOTYPE (wild-type, Large^+/−^) and SEX (male, female). For the wild-type placenta PWS gene (*Snhg14* or *Ndn*) molecule count data, a Mann-Whitney-Wilcoxon test was used to determine if there was a significant difference between cell type (either *Kdr*+ [endothelial] or *Kdr*– [non-endothelial]) due to the fact the data was non-parametric and unpaired. All statistical tests for this step were performed in RStudio and an P value of 0.05 was used, before correction for multiple testing using the Bonferroni method.

### MicroCT

Six Large^+/−^ and six wild-type embryos (three male, three female for each genotype, derived from two separate litters) were dissected into ice cold phosphate buffered saline (PBS) and exsanguinated by severing the umbilical vessels. After washing in PBS, embryos were fixed by immersion in 4% paraformaldehyde (PFA in PBS) for 7 days at 4°C, before storing in 1% PFA at 4°C.

Embryos were contrasted by immersion in 50% Lugol’s solution (1:1 Lugold:ddH2)) for 2-weeks, protected from light, and solution exchanged for fresh every 2-days. After contrasting, samples were washed in ddH2O for at least 1-hour before embedding in an acrylic mount in 1% agarose dissolved in ddH2O and allowed to set for at least 2-hours.

MicroCT data sets were acquired using a Skyscan 1272 scanner (Bruker) with the x-ray source set to 70kV and using 1mm aluminium filter. Pixel resolution of 15µm/pixel was set at 3×3 camera pixel binning. Four projections were acquired and averaged every 0.4° through a total rotation of 360° With a final step of 25 random movements. NRecon (Bruker) was used for 3D reconstruction to produce a final volume in a PNG image-stack format.

For image analysis, each MicroCT dataset was imported into the commercial software Avizo (Avizo 3D 2024.2, ThermoFisher). The whole embryo, brain, and kidney were manually segmented using Avizo’s segmentation editor. The organ volume for each embryo was normalised by total embryo volume to enable a direct comparison. The average intensity (note the tissue was iodinated) of the organs was measured by masking the organs of the 16bit depth volume and summing all the pixel intensities in the mask divided by the volume of the mask. The segmented organ volume was used to mask eah organ from the total embryo.

Although both males and females were included in the sampling we were under-powered to detect GENOTYPE*SEX interactions and so chose to use ANCOVA to analyse the data, with GENOTYPE as a fixed factor and SEX as a co-factor. All statistical tests for this step were performed in SPSS and an P value of 0.05 was used, before correction for multiple testing using the Bonferroni method.

## Acknowledgements

The authors thank the Molecular and Cellular Biology Team at the MRC Mary Lyon Centre, Oxfordshire for their technical support in generating *Large*^+/−^ samples. The authors also thank Dr Theresa Strong of the Foundation for Prader-Willi Research for helpful discussions relating to this project.

## Competing interests

The authors have no competing interests.

## Funding

This work was supported by a grant to ARI and RMJ from the Foundation for Prader-Willi Research (FPWR, https://ror.org/05dxwnm86) and by the FPWR Pre-Clinical Animal Network. X-ray CT image analysis was undertaken by the National Research Facility for Lab X-ray CT (NXCT; nxct.ac.uk) at the µ-VIS X-ray Imaging Centre (muvis.org), University of Southampton, funded through EPSRC grant EP/T02593X/1. This was also supported by an NVIDIA grant and utilised an NVIDIA RTX A6000.

## Data and resource availability

All details of resources and statistical analyses can be found within the article and its supplementary information. All QC’d data are publicly available via ‘Open science framework’: doi 10.17605/OSF.IO/B2PDV

## Author contributions

AW – Data Curation, Formal Analysis, Investigation, Validation, Visualization, Writing (Reviewing and Editing); Raquel BS – Investigation, Methodology, Visualization; LB – Investigation, Data Curation, Formal Analysis, Writing (Reviewing and Editing); CC – Investigation, Data Curation, Formal Analysis, Writing (Reviewing and Editing); RAJ – Formal Analysis; Rosie BS – Investigation; MS – Supervision; SEW – Supervision; DSC – Data Curation, Formal analysis; SKR – Funding Acquisition, Supervision; MJH – Methodology, Supervision; RMJ – Conceptualization, Funding Acquisition, Supervision, Writing (Reviewing and Editing); ARI – Conceptualization, Funding Acquisition, Project Administration, Supervision, Writing (Original Draft Preparation).

## Notes

### Competing Interest Statement

The authors have declared no competing interest.

### Summary of Updates

Updated Data and Resource availability section including doi for data.

