## Supporting Information for "Placental defects revealed by modelling PWS in mice"

### Placental defects revealed by modelling PWS in mice – SUPPORTING INFORMATION

#### SUPPORTING RESULTS

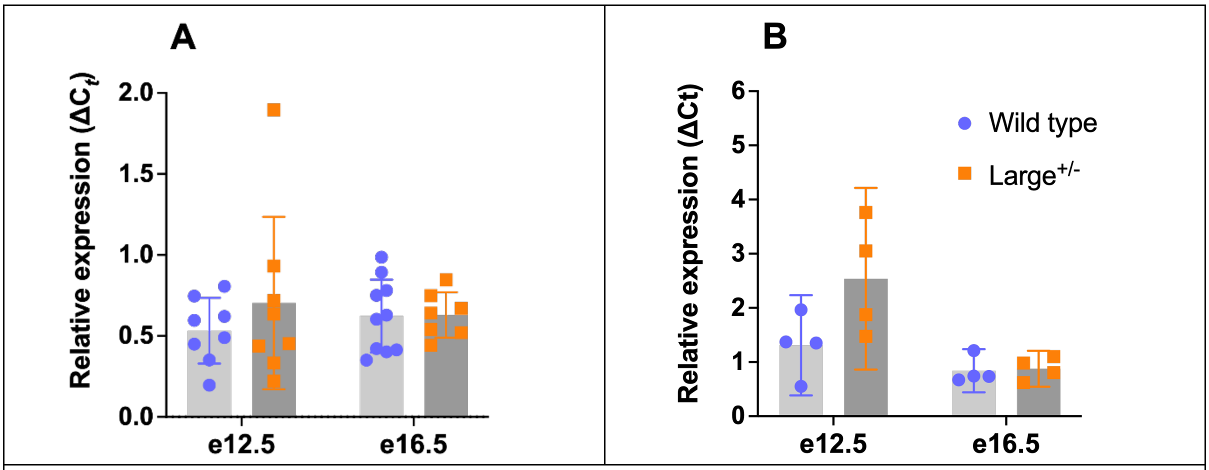


**Figure S1** Expression of *Ube3a* in placenta (**A**) and whole brain (**B**) homogenate from wild-type and Large$^{+/-}$ mice at two different embryonic timepoints. Gene expression normalised to the geometric mean of three house-keeping genes using the ΔCt method. Bars are means with 95% CI indicated.

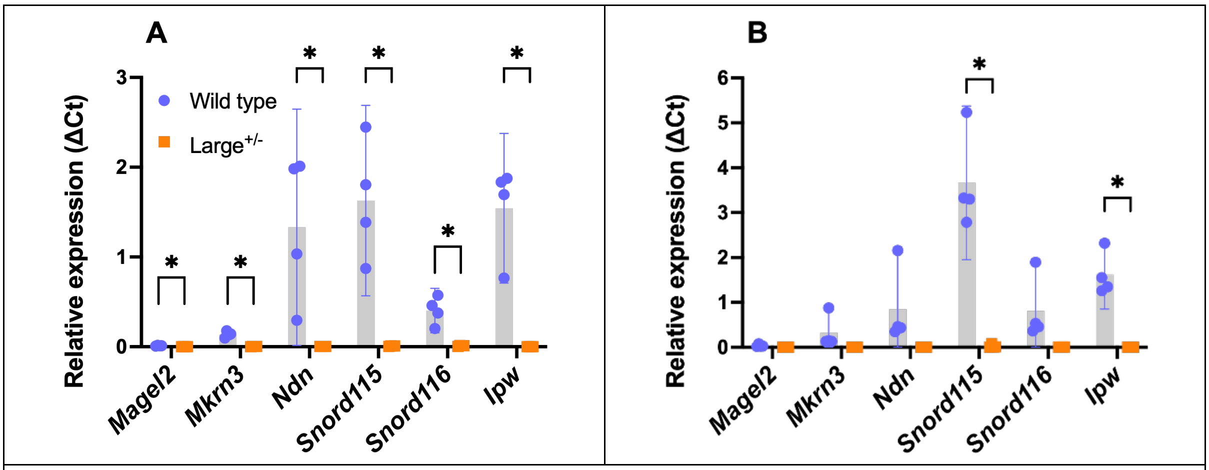


**Figure S2** Expression of PWS genes in whole brain homogenate from wild-type and Large$^{+/-}$ mice at two different embryonic timepoints. At e12.5 (**A**), expression of all PWS genes in the placenta was significantly reduced in Large$^{+/-}$ mice relative to wild-type controls. At e16.5 (**B**), although means expression was reduced in Large$^{+/-}$ mice, this only reached statistical significance for two PWS genes. Gene expression normalised to the geometric mean of three house-keeping genes using the ΔCt method. Bars are means with 95% CI indicated.

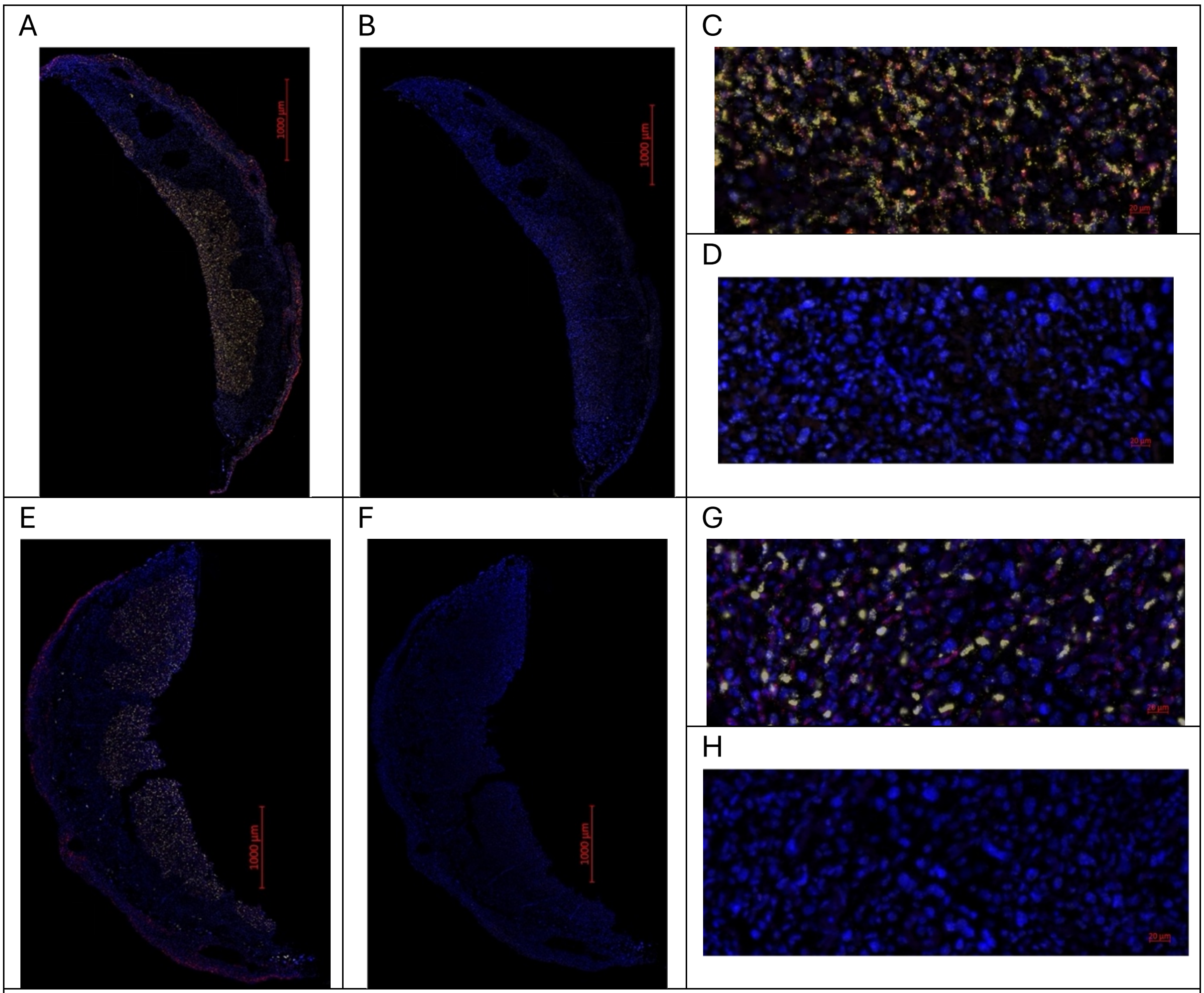


**Figure S3**. Representative images of RNAscope analysis of *Ndn* (**A** and **C**) and *Snhg14* (**E** and **G**) and the endothelial cell marker *Kdr* in wild type placenta. No-probe control images for *Ndn/Kdr* (**B** and **D**) and *Snhg14/Kdr* (**F** and **H**) illustrate the level of background fluorescence for each sample that was accounted for in the image analysis.

| | Factors investigated via two-way ANOVA test | | | | | |
|---|---|---|---|---|---|---|
| | **SEX** | | **TIMEPOINT** | | **SEX*TIMEPOINT** | |
| **Gene name** | **F value** | **P-value** | **F value** | **P-value** | **F value** | **P-value** |
| *Mkrn3* | 1.44 | 0.25 | 1.51 | 0.25 | 0.97 | 0.40 |
| *Ube3a* | 0.23 | 0.64 | 5.67 | **0.012** | 0.48 | 0.62 |
| *Ipw* | 1.96 | 0.18 | 3.93 | **0.04** | 2.61 | 0.10 |
| *Snord115* | 0.002 | 0.97 | 5.83 | **0.011** | 1.47 | 0.26 |
| *Snord116* | 0.14 | 0.71 | 3.07 | **0.07** | 1.84 | 0.19 |

**Table S1** Statistics for two-way ANOVA analysis of PWS gene expression in wild type placenta across three embryonic timepoints (e10.5, e12.5 and e16.5). Based on the results of the Shapiro-Wilk test, normally distributed data were analysed by ANOVA using the factors TIMEPOINT and SEX. Statistically significant differences are highlighted in bold.

| **Gene name** | **Kruskal-Wallis test;** TIMEPOINT | | **Kruskal effect size value** | **Kruskal-Wallis test;** SEX | | **Kruskal effect size value** |
|---|---|---|---|---|---|---|
| | **$\chi^2$ statistic** | **P-value** | | **$\chi^2$ statistic** | **P-value** | |
| *Magel2* | 15.86 | **0.0004** | 0.66 | 0.003 | 0.954 | -0.045 |
| *Ndn* | 8.66 | **0.013** | 0.317 | 1.47 | 0.225 | 0.021 |

**Table S2** Statistics for Kruskal-Wallis analysis of wild-type placental PWS gene expression in wild type placenta across three embryonic timepoints (e10.5, e12.5 and e16.5). Non-normally distributed data as determined by the Shapiro-Wilk test, were analysed by Kruskal-Wallis using the factors TIMEPOINT and SEX. Statistically significant differences are highlighted in bold.

| **Gene name** | **E10.5 – E12.5** | | **E10.5 – E16.5** | | **E12.5 – E16.5** | |
|---|---|---|---|---|---|---|
| | **Z statistic** | **Adjusted p-value** | **Z statistic** | **Adjusted p-value** | **Z statistic** | **Adjusted p-value** |
| *Mkrn3* | -0.88 | 1 | 0.57 | 1 | 1.45 | 0.44 |
| *Magel2* | -3.75 | **0.0005** | -3.04 | **0.007** | 0.707 | 1 |
| *Ndn* | -0.85 | 1 | 2.02 | 0.1 | 2.86 | **0.004** |
| *Snord116* | -0.92 | 1 | 1.66 | 0.29 | 2.58 | **0.03** |
| *Ipw* | 2.19 | 0.09 | 1.2 | 0.7 | -0.99 | 1 |
| *Snord115* | -1.03 | 0.9 | 1.98 | 0.1 | 3.01 | **0.008** |

**Table S3** Results of *post-hoc* Dunn pairwise tests on wild type PWS genes placental gene expression across three embryonic timepoints, including the z statistic and the Bonferroni corrected p-value. Statistically significant differences are highlighted in bold.

| | **Factors investigated via two-way ANOVA test** | | | | | |
|---|---|---|---|---|---|---|
| | **GENOTYPE** | | **SEX** | | **GENOTYPE*SEX** | |
| **Gene name** | **F value** | **P-value** | **F value** | **P-value** | **F value** | **P-value** |
| *Magel2* | 91.16 | **5.89E-07** | 0.49 | 0.498 | 3.46 | 0.088 |
| *Snord115* | 59.19 | **1.17E-04** | 0.70 | 0.695 | 1.17 | 0.314 |
| *Ipw* | 14.70 | **0.0024** | 0.29 | 0.287 | 0.03 | 0.858 |

**Table S4** Statistics for two-way ANOVA analysis of PWS gene expression in wild type and Large$^{+/-}$ placenta at e12.5. Based on the results of the Shapiro-Wilk test, normally distributed data were analysed by ANOVA using the factors GENOTYPE and SEX. Statistically significant differences are highlighted in bold.

| **Gene name** | **Kruskal-Wallis test;** GENOTYPE | | **Kruskal effect size value** | **Kruskal-Wallis test;** SEX | | **Kruskal effect size value** |
|---|---|---|---|---|---|---|
| | **χ2 statistic** | **P-value** | | **χ2 statistic** | **P-value** | |
| *Mkrn3* | 7.5 | **0.0063** | 0.461 | 0.10 | 0.753 | -0.064 |
| *Ndn* | 10.5 | **0.0012** | 0.679 | 0.05 | 0.817 | -0.068 |
| *Snord116* | 7.5 | **0.0062** | 0.464 | 2.70 | 0.100 | 0.121 |

**Table S5** Statistics for Kruskal-Wallis analysis of wild-type placental PWS gene expression in wild type and Large$^{+/-}$ placenta at e12.5. Non-normally distributed data as determined by the Shapiro-Wilk test, were analysed by Kruskal-Wallis using the factors GENOTYPE and SEX. Statistically significant differences are highlighted in bold.

| | Factors investigated via two-way ANOVA test | | | | | |
|---|---|---|---|---|---|---|
| | **GENOTYPE** | | **SEX** | | **GENOTYPE*SEX** | |
| **Gene name** | **F value** | **P-value** | **F value** | **P-value** | **F value** | **P-value** |
| *Magel2* | 62.698 | **2.5E-06** | 0.839 | 0.376 | 0.789 | 0.391 |
| *Snord115* | 4.511 | 0.053 | 0.200 | 0.662 | 1.892 | 0.192 |
| *Snord116* | 0.936 | 0.351 | 0.263 | 0.167 | 1.520 | 0.239 |
| *Ndn* | 215.593 | **1.8E-09** | 0.146 | 0.708 | 0.340 | 0.570 |
| *Mkrn3* | 2.734 | 0.122 | 0.062 | 0.808 | 0.001 | 0.979 |
| *Ipw* | 57.387 | **4.0E-06** | 0.636 | 0.439 | 0.016 | 0.900 |

**Table S6** Statistics for two-way ANOVA analysis of PWS gene expression in wild type and Large$^{+/-}$ placenta at e16.5. Based on the results of the Shapiro-Wilk test, normally distributed data were analysed by ANOVA using the factors GENOTYPE and SEX. Statistically significant differences are highlighted in bold.

| | | Average total cell counts (±SD) | Average *Kdr*-positive cell counts (±SD) |
|---|---|---|---|
| Wild type placenta | Male | 23,259 (5,885) | 10,040 (687) |
| | Female | 23,323 (4,400) | 8,448 (2807 |
| Large$^{+/-}$ placenta | Male | 22,918 (5,048) | 4846 (3436) |
| | Female | 22,207 (5,973) | 4835 (754) |

**Table S7** Total cell counts for placenta (DAPI stained cells) and endothelial cell (*Kdr*-positive) broken down by genotype and sex. The individual counts were used to derive the percentage *Kdr*-positive cell values used in Figure 4G of the Main text.

#### SUPPORTING METHODS

| Gene Primer | | 5'-3' Sequence | Optimum Conc. (nM) |
|---|---|---|---|
| *Mrkn3* | For | CCAATCAGTTGCTTAAGAAGTTGC | 300 |
| | Rev | AAGAGCCAACGGTCATCAGAG | 300 |
| *Magel2* | For | GCATAGCAAGCCAGCCTCAG | 300 |
| | Rev | GTAGACGAGCCTGTGGAGCCT | 50 |
| *Necdin* (*Ndn*) | For | ATGGTGCAGAAGCATCCTCAG | 300 |
| | Rev | ATGGTGTGGAGATTGGTCAGC | 300 |
| *Snord116* | For | TGGATCTATGATGATTCCCAG | 300 |
| | Rev | TGGACCTCAGTTCCGATGAG | 50 |
| *Ipw* | For | TGCTGTTATTGCCTTGCCTG | 300 |
| | Rev | TGGTGAAGCTGCTGGTAGAA | 300 |
| *Snord115* | For | ACAACCCACTGTCATGAAGAAAGG | 50 |
| | Rev | CCTCAGCGTAATCCTATTGAGCAT | 700 |
| *Ube3a* | For | CAGACGTGACCATATTATAGATGATGC | 300 |
| | Rev | CCACATACAACTGCTTCTTCAAGTCT | 300 |
| *Hprt* | For | GCGATGATGAACCAGGTTATGA | 300 |
| | Rev | GCCTCCCATCTCCTTCATGA | 300 |
| *Dynein* | For | GACCTCAGGCTCAGACGAAGAC | 300 |
| | Rev | AAGACGCTCATGGCATCACA | 300 |
| *Beta-actin* | For | TTCTGGTGCTTGTCTCACTGA | 300 |
| | Rev | CAGTATGTTCGGCTTCCCATTC | 300 |

**Table S7** Primer sequences and concentrations used in qPCR
